# Young adults who improve performance during dual-task walking show more flexible reallocation of cognitive resources: A Mobile Brain-Body Imaging (MoBI) study

**DOI:** 10.1101/2022.03.18.484948

**Authors:** Eleni Patelaki, John J. Foxe, Kevin A. Mazurek, Edward G. Freedman

**Affiliations:** The Frederick J. and Marion A. Schindler Cognitive Neurophysiology Laboratory, The Del Monte Institute for Neuroscience, Department of Neuroscience, University of Rochester School of Medicine and Dentistry, Rochester, New York, USA; Department of Biomedical Engineering, University of Rochester, Rochester, New York, USA; Department of Physiology and Biomedical Engineering, Mayo Clinic, Rochester, Minnesota, USA; The Well Living Lab, Rochester, Minnesota, USA

**Keywords:** Event-related potentials, EEG, Response inhibition, Behavior, Gait, Error processing, Biological Sciences – Neuroscience

## Abstract

In young adults, pairing a taxing cognitive task with walking can have different effects on gait and cognitive task performance. In some cases, performance clearly declines whereas in others compensatory mechanisms maintain performance even under dual-task conditions. This study set out to investigate the preliminary finding of behavioral improvement in Go-NoGo response inhibition task performance during walking compared to sitting, which was observed at the piloting stage. Mobile Brain/Body Imaging (MoBI) was used to record electroencephalographic (EEG) activity, three-dimensional (3D) gait kinematics and behavioral responses in the cognitive task, during sitting or walking on a treadmill. In a cohort of twenty-six (26) young adults, fourteen (14) participants improved in measures of cognitive task performance while walking compared to sitting. These participants exhibited walking-related EEG amplitude reductions over frontal brain scalp regions during key stages of inhibitory control (conflict monitoring, control implementation and pre-motor stages), accompanied by reduced stride-to-stride variability and faster responses to stimuli compared to those who did not improve. In contrast, the twelve (12) participants who did not improve exhibited no EEG amplitude differences across physical condition. The neural activity changes associated with performance improvement during dual tasking hold promise as cognitive flexibility markers that can potentially help assess cognitive decline in aging and neurodegeneration.

## Introduction

Performance of executive functions requires coordination across distributed neural networks for both routine and more complex cognitive processes [1-4]. Yet, there are limits to the number and complexity of tasks that can be undertaken at the same time [5]. As this capacity limit is approached, behavioral performance may begin to deteriorate [6, 7]. Dual task performance, the simultaneous performance of two tasks, activates multiple brain regions concurrently, which can tax cognitive systems and bring them closer to this capacity limit [8-10]. Performing cognitive and motor tasks simultaneously sets the stage for competition for available neural resources leading to performance declines in both modalities. This is referred to as cognitive-motor interference (CMI) [11-14]. Pairing cognitive tasks with walking can elicit dual-task decline in gait performance and task-related behavior, as well as altered patterns of neural activation in older neurotypical adults [15-22] and in various patient populations [23-28]. However, in young adults, the manifestation of decrements in gait and cognitive task performance is not as clear, and as such, the neural activity changes detected in this group reflect interaction but not necessarily interference at a neural resource level. Some studies have reported evidence of no deterioration of response accuracy or increases in gait variability during dual-task walking in young adults [18, 29-36]. These studies suggest that young healthy adults adapt their gait and task-related behavior during dual-task walking and, consistent with this conclusion, report slower reaction times to stimuli [30-32], changes in stride length [29, 33, 36] and reduced gait speed and velocity [18, 33, 35]. These findings indicate that young adults adopt a more deliberate approach to both task responses and walking in order to maintain task accuracy, and as such, point to strategy changes that will necessarily involve neural reconfigurations. On the other hand, there are studies that have reported reductions in response accuracy [37, 38] and increases in gait variability [39-41] during dual-task walking in young adults. This discrepancy suggests that young adults do not reach their cognitive capacity limit under all dual-task walking conditions; under certain conditions, presumably when the dual-task load is below the capacity limit, they seem to activate mechanisms to compensate for the increased dual-task demands. In these cases, dual-task-related neural activity changes likely reflect a reallocation of neural resources and the adoption of a different cognitive strategy that drives the observed compensatory adaptations.

Response inhibition, namely withholding response to a thought, emotion, or stimulus, is one of the core executive functions and a vital component of everyday living. One often-used approach to studying response inhibition is the Go/NoGo task using a set of visual images as stimuli. The task requires pressing a response button after each novel image is presented (‘Go’ trial), but withholding the button press in response to the second presentation of a repeated image (‘NoGo’ trial) [15, 23, 29, 42-48]. During successful NoGo trials during which the participant properly withholds a response, two stimulus-locked Event-Related Potential (ERP) components are typically elicited: the N2 and the P3. The N2 is a negative voltage deflection that peaks around 200-350 ms [49, 50] post-stimulus-onset and has a frontocentral scalp distribution. This topographical distribution reflects its generation by the anterior cingulate cortex (ACC), a brain region which is key for monitoring inhibitory conflict [49, 51-53]. The P3 is a positive voltage deflection that peaks around 350-600 ms post-stimulus-onset and has a broad distribution, extending from centroparietal to frontal areas [29, 54]. During the P3 processing stage, both motor and cognitive components of inhibition are executed [47]. In parallel to the motor inhibitory component of button press cancellation, higher inhibitory control is putatively exerted by lateral prefrontal areas, specifically the dorsolateral prefrontal cortex (DLPFC), to reduce inhibitory conflict in ensuing trials and hence improve cognitive task performance [55, 56]. During unsuccessful NoGo inhibition trials, an ERP component known as the response-locked Event-Related Negativity (ERN) is elicited. The frontocentral ERN peaks approximately 50 ms after the erroneous motor response [57-59], and reflects conflict monitoring in error trials and the source has been localized to the ACC [55, 60, 61].

The effects of walking on the N2 and P3 typically elicited during successful performance of the Go-NoGo response inhibition task has been investigated by previous studies [15, 29]. De Sanctis and colleagues [29] observed amplitude reductions of both the N2 and the P3 event-related potentials (ERPs) during successful inhibitions while walking, as well as an anteriorization of the P3 distribution suggesting recruitment of frontal cortical circuits. Furthermore, they found no significant differences between sitting and walking in terms of response accuracy and response speed, and no significant changes in stride-to-stride variability when comparing single-task and dual-task walking in young adults. In the absence of significant dual-task decrements, these findings were interpreted as a shift to a less automatic (N2 amplitude reduction) and more effortful (P3 frontalization) cognitive strategy during walking.

Other studies focused on the correlation between neural activity and several behavioral measures in the context of a Go-NoGo response inhibition task. Falkenstein and colleagues found that young healthy individuals who had a high rate of unsuccessful inhibitions exhibited a smaller and later N2 compared to those with a low rate of unsuccessful inhibitions [62]. Roche and colleagues showed that highly absentminded, young healthy individuals had larger and earlier N2 and P3 components in successful inhibition trials and larger error-related components in unsuccessful inhibition trials compared to less absentminded individuals of the same age group [63]. Karamacoska and colleagues reported smaller P3 amplitudes in young healthy adults with increased response time variability, who were also found to commit more errors, compared to peers with low response time variability [64]. In each case, different criteria were used to split the cohort into two subgroups, depending on the behavioral variable of interest: certain studies performed a median split [46, 63-65], others leveraged the bimodality of the distribution of the behavioral variable and split the cohort based on the two modes [62] and studies investigating impulsivity applied the splitting methodology proposed by Pailing and colleagues [66, 67]. In the context of collecting pilot data for the present study, data from five (5) young healthy adults were collected. Analysis of these preliminary data showed that three (3) of the pilot participants improved response accuracy during walking compared to sitting. Improvement in cognitive task performance with the addition of walking appeared to conflict with the CMI hypothesis. Response accuracy was measured using the d’ score (sensitivity index) [68, 69], since it is a bias-free measure that takes into account both the Go and the NoGo behavior (greater d’ score signifies better response accuracy). These preliminary data were then sequestered (see Supplemental Material). The working hypothesis that young individuals who improve performance during walking would differ in their ability to flexibly allocate neural resources for accomplishing both the motoric and cognitive tasks, and might also differ in the consistency of their gait compared to those who do not improve while walking, was tested in this report. Participants were divided into two (2) subgroups based on the walking-*minus*-sitting d’ difference: 1) participants who exhibited a positive walking-*minus*-sitting d’ difference (cognitive task performance improved during walking compared to sitting - **IMPs**) and 2) participants who exhibited a walking-*minus*-sitting d’ difference which was either negative or not significantly different from zero (cognitive task performance did not improve during walking compared to sitting - **nIMPs**).

The current study examined successful NoGo inhibition trials during the N2 and P3 processing stages and unsuccessful NoGo inhibition trials during the ERN stage for walking-related amplitude changes in neurophysiological activity in the entire young adult cohort, and subsequently in the IMP and nIMP subgroups. Gait variability and response speed were additionally examined for dual-task changes in the same groups, to test whether improvement in response accuracy (IMPs) would be accompanied by trade-offs reflected in other physiological domains; for example, whether IMPs would be more accurate but slower in their responses, or they would walk more variably. Identifying potential differences in the way IMPs alter their neural activity in response to dual-task load compared to nIMPs can shed light on the underlying neurocognitive mechanisms that drive their behavioral improvement.

## Materials and Methods

### Participants

Twenty-six (26) young adults (18-30 years old; age = 22.35 ± 3.27 years; 13 female, 13 male; 23 right-handed, 3 left-handed) participated in the study. All participants provided written informed consent, reported no diagnosed neurological conditions, no recent head injuries, and normal or corrected-to-normal vision. The Institutional Review Board of the University of Rochester approved the experimental procedures (STUDY00001952). All procedures were compliant with the principles laid out in the Declaration of Helsinki for the responsible conduct of research. Participants were paid $15/hour for time spent in the lab.

### Experimental Design

A Go-NoGo response inhibition cognitive task was employed. During each experimental block, images were presented in the central visual field for 67 ms with a fixed stimulus-onset-asynchrony of 1017 ms. Images subtended 10° of visual angle horizontally and 8° vertically. The task was coded using the Presentation software (version 20.1, Neurobehavioral Systems, Albany, CA, USA). Participants were instructed to press the button of a wireless game controller as fast and accurately as possible if the presented image was different from the preceding image (‘Go’ trial). They were instructed to withhold pressing the button if the presented image was the same as the preceding image (‘NoGo’ trial) (Fig. 1). Participants performed blocks of 240 trials in which 209 (87%) were Go trials and 31 (13%) were NoGo trials. NoGo trials were randomly distributed within each block.

**Fig. 1.**
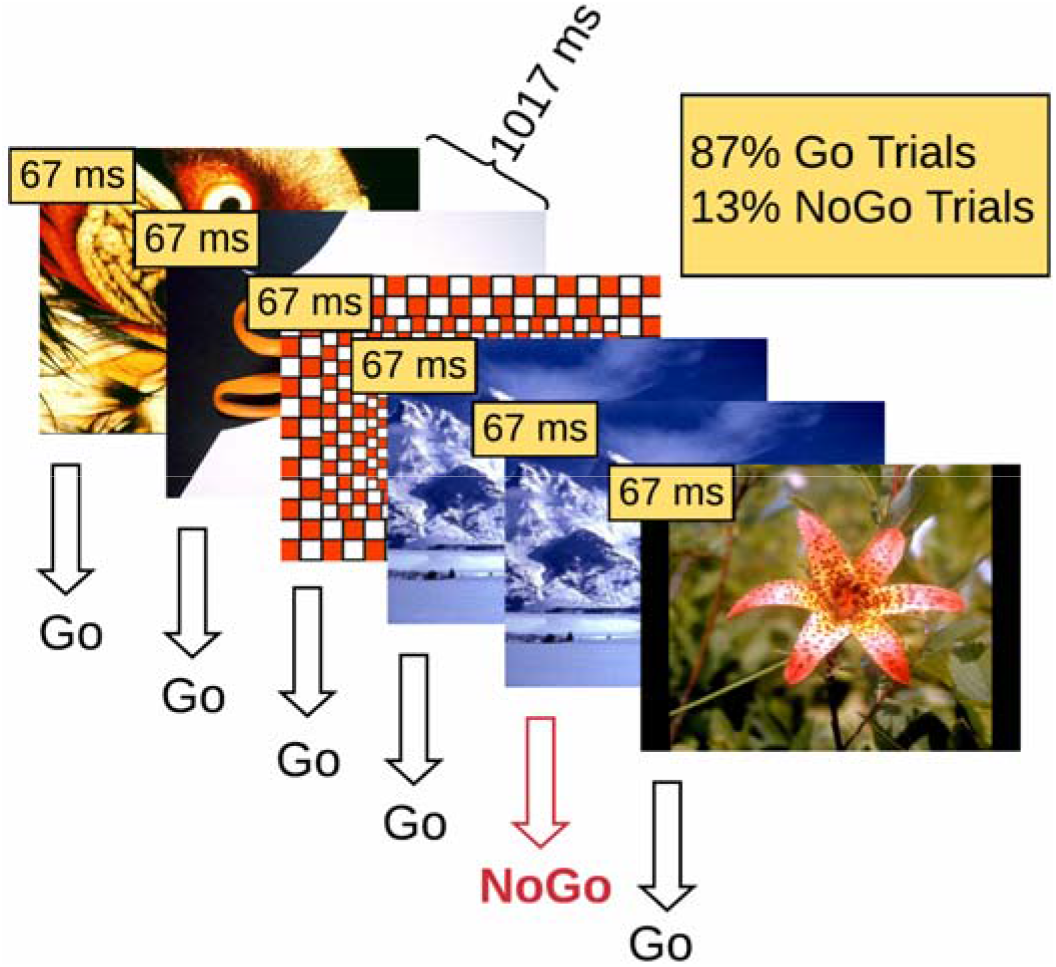
Illustration of the Go-NoGo response inhibition experimental design. Participants are instructed to respond on Go trials and withhold response on NoGo trials.

Three (3) behavioral conditions of the cognitive task were defined: 1) correct rejections, defined as the NoGo trials on which participants correctly withheld their response, 2) false alarms, defined as the NoGo trials on which participants incorrectly pressed the response button and 3) hits, defined as the Go trials on which participants correctly pressed the response button.

Experimental blocks were performed while the participants were either sitting or walking on a treadmill (Tuff Tread, Conroe, TX, USA), at a distance of 2.25 m approximately from the projection screen on which the images were projected (Barco F35 AS3D, 1920×1080 pxl). A safety harness was worn while walking to guard against falls [70]. An experimental session consisted of 16 blocks: 1 training block at the beginning, 7 sitting blocks, 7 walking blocks, and 1 single-task walking block (walking on the treadmill without a cognitive task). The order of sitting and walking blocks was pseudorandomized; no more than 3 consecutive walking blocks occurred to prevent exhausting the participants. Participants were allowed to take short breaks between the blocks, each of which lasted 4 minutes. Most participants took at least one break during the experiment. If a break was requested, typically it did not last longer than 10 minutes. Participants were asked to select a treadmill speed corresponding to brisk walking for them, starting from the recommended speed of 4.8 km/h and increasing or decreasing as necessary. The vast majority of participants (22 out of 26) selected a speed of 4.8 km/h, while 4 participants selected lower speeds (3 participants walked at 4.2 km/h and 1 participant at 3.9 km/h). In general, the walking speeds selected corresponded to brisk walking [71].

The pictures used for stimuli were drawn from the International Affective Picture System (IAPS) database [72]. The IAPS database contains pictures of varied emotional valence and semantic content. Positive, neutral and negative pictures were all used, however analyzing the emotional valence or semantic content of stimuli is beyond the scope of this study.

EEG data were recorded using a BioSemi Active Two System (BioSemi Inc., Amsterdam, The Netherlands) and a 64-electrode configuration following the International 10-20 system. Neural activity was digitized at 2048 Hz. Full-body motion capture was recorded using a 16 camera OptiTrack system (Prime 41 cameras), and Motive software (OptiTrack, NaturalPoint, Inc., Corvallis, OR, USA) in a ~37 m^2^ space. Cameras recorded 41 markers on standard anatomical landmarks along the torso, the head and both arms, hands, legs and feet at 360 frames per second. Stimulus triggers from Presentation (Neurobehavioral Systems Inc., Berkeley, CA, USA), behavioral responses from the game controller button, motion tracking data and EEG data were time-synchronized using Lab Streaming Layer (LSL) software (Swartz Center for Computational Neuroscience, University of California, San Diego, CA, USA; available at: https://github.com/sccn/labstreaminglayer). Motion capture data were recorded using custom software written to rebroadcast the data from the Motive software to the LSL lab recorder. EEG data were recorded from available LSL streaming plugins for the BioSemi system. Behavioral event markers were recorded using the built-in LSL functionality in the Presentation software. The long-term test-retest reliability of the MoBI approach has been recently detailed [73]. All behavioral, EEG and motion kinematic data processing and basic analyses were performed using custom MATLAB scripts (MathWorks Inc., Natick, MA, USA) and/or functions from EEGLAB [74]. Custom code from this study will be made available on GitHub (https://github.com/CNL-R) upon publication.

### Cognitive Task Performance Processing & Analysis

The exact timing of each button press relative to stimulus onset, the participant’s response times (RTs), were recorded using the Response Manager functionality of Presentation and stored with precision of 1/10 millisecond. The Response Manager was set to accept responses only after 183 ms post-stimulus-onset within each experimental trial. Any responses prior to that were considered delayed responses to the previous trial and were ignored. This RT threshold was selected to filter out as many delayed-response trials as possible, without rejecting any valid trials for which the responses were merely fast [75].

Two (2) behavioral conditions of the cognitive task were interrogated in this study, 1) correct rejections and 2) false alarms. For both correct rejections and false alarms, only trials that were preceded by hits were kept, to ensure that the inhibitory component was present.

Two (2) behavioral measures were calculated: 1) the d’ score (sensitivity index) and 2) mean RT during (correct) Go trials, namely hits. D’ is a standardized score and it is computed as the difference between the Gaussian standard scores for the false alarm rate and the hit rate [68, 69].

#### Statistical Analysis

In the full young adult cohort, d’ score differences between sitting and walking were assessed using a paired t-test (the walking-*minus*-sitting d’ score difference was subjected to a Shapiro-Wilk normality test [76] and the null hypothesis was not rejected). Additionally, mean RTs during Go trials were tested for differences between sitting and walking using a paired t-test (the walking-*minus*-sitting mean RT difference was subjected to a Shapiro-Wilk normality test and the null hypothesis was not rejected).

Participants were subsequently classified on the basis of whether their d’ score during walking was significantly higher than when they were seated (their cognitive task performance improved **(IMP)** while walking), or whether they did not improve (**nIMP**) performance while walking (either because their d’ scores declined or were unchanged). Significant walking-*minus*-sitting d’ score difference was defined as the difference that lay outside of the 95% confidence interval of the normal distribution that had a mean value of zero and a standard deviation equal to that of the (d’walking – d’sitting) distribution of the entire cohort.

In the context of the split-group analysis, IMPs and nIMPs were compared in terms of average d’ score, i.e. 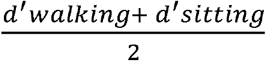, using an independent samples t-test (the average d’ scores of IMPs and IMPs were subjected to a Shapiro-Wilk normality test and the null hypothesis was not rejected for either group). Additionally, mean RTs during Go trials were subjected to a 2 (Group: IMPs, nIMPs) × 2 (Motor Load: sitting, walking) ANOVA to test for response speed differences between IMPs and nIMPs, as well as for differences in how response speed was modulated by the addition of walking in these groups.

### Gait Kinematics Processing & Analysis

Heel markers on each foot were used to track gait kinematics. The three dimensions (3D) of movement were defined as follows: X is the dimension of lateral movement (right-and-left relative to the motion of the treadmill belt), Y is the dimension of vertical movement (up-and-down relative to the motion of the treadmill belt), and Z is the dimension of fore-aft movement (parallel to the motion of the treadmill belt). The heel marker motion in 3D is described by the three (3) time series of the marker position over time in the X, Y and Z dimension, respectively. Gait cycle was defined as the time interval between two consecutive heel strikes of the same foot. Heel strikes were identified as the local maxima of the Z position waveform over time. To ensure that no ‘phantom’ heel strikes were captured, only peaks with a prominence greater than 0.1 m were kept (*findpeaks* function in MATLAB, *minimum peak prominence* parameter was set to 0.1 m).

Stride-to-stride variability was quantified as the mean Euclidean distance between consecutive 3D gait cycle trajectories, using the Dynamic Time Warping algorithm (DTW) [77, 78]. DTW is an algorithm for measuring the similarity between time series, and its efficacy in measuring 3D gait trajectory similarity is well-established [79-81].

In the case of one-dimensional signals, if X_m=1,2,..,M_ the reference signal and Y_n=1,2,..,N_ the test signal, then DTW finds a sequence {ix, iy} of indices (called warping path), such that X(ix) and Y(iy) have the smallest possible distance. The ix and iy are monotonically increasing indices to the elements of signals X, Y respectively, such that elements of these signals can be indexed repeatedly as many times as necessary to expand appropriate portions of the signals and thus achieve the optimal match. This concept can be generalized to multidimensional signals too, like the 3D gait cycle trajectories which are of interest here. The minimal distance between the reference and the test signals (gait trajectories here) is given by the equation below:

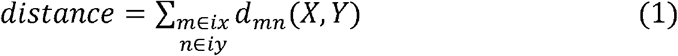

Gait cycle trajectories with a kurtosis that exceeded 5 standard deviations of the mean were rejected as outliers. Also, before DTW computation, gait cycle trajectories were resampled to 100 samples. Since DTW essentially calculates the sum of the Euclidean distances between corresponding points of two interrogated trajectories, ensuring that all trajectories are resampled to the same length helps avoid bias in the algorithm computations.

The actual measure that was used to quantify each participant’s stride-to-stride variability is the mean across DTW distances occurring from all stride-to-stride comparisons. Right-foot and left-foot stride-to-stride DTW distances were pooled to calculate the mean DTW distance per participant.

#### Statistical Analysis

In the full young adult cohort, mean DTW distance differences between single-task (ST) walking and dual-task (DT) walking were assessed using a Wilcoxon signed rank test (the DT-*minus*-ST mean DTW distance difference was subjected to a Shapiro-Wilk normality test and the null hypothesis was rejected). One (1) participant did not have ST walking recordings and was therefore excluded from this analysis.

In the context of the split-group analysis, mean DTW distance was subjected to a 2 (Group: IMPs, nIMPs) x 2 (Cognitive Load: ST walking, DT walking) ANOVA to test for stride-to-stride variability differences between IMPs and nIMPs, as well as for differences in how the addition of cognitive task performance modulated stride-to-stride variability in these groups. The two participants excluded from the group level analysis both belonged to the IMP subgroup.

### EEG Activity Processing & Analysis

EEG signals were first filtered using a zero-phase Chebyshev Type II filter (*filtfilt* function in MATLAB, passband ripple *Apass* = 1 dB, stopband attenuation *Astop* = 65 dB) [82], and subsequently down-sampled from 2048 Hz to 512 Hz. Next, ‘bad’ electrodes were detected based on kurtosis, probability, and spectrum of the recorded data, setting the threshold to 5 standard deviations of the mean, as well as covariance, with the threshold set to ±3 standard deviations of the mean [82]. These ‘bad’ electrodes were removed and interpolated based on neighboring electrodes, using spherical interpolation. All the electrodes were re-referenced offline to a common average reference.

Winkler and colleagues have shown that 1-2 Hz highpass filtered EEG data yield the optimal Independent Component Analysis (ICA) decomposition results in terms of signal-to-noise ratio [83]. In order to both achieve a high-quality ICA decomposition and retain as much low-frequency (< 1 Hz) neural activity as possible, after running Infomax ICA (*runica* function in EEGLAB, the number of retained principal components matched the rank of the EEG data) on 1-45 Hz bandpass-filtered data and obtaining the decomposition matrices (weight and sphere matrices), these matrices were transferred and applied to 45-Hz lowpass-filtered data. No highpass filtering was applied, since there is evidence indicating that the best way to avoid introducing artifacts into the ERP waveforms is either to use conservative high-pass filters (≤ 0.1 Hz) or to avoid high-pass filtering altogether [84]. ICs were labeled using the ICLabel algorithm [85]. Artifactual ICs, namely ICs classified as eye activity, muscle activity, ground noise, poor electrode quality or contact, or heart activity, were detected and rejected, while the remaining ICs were back-projected to the sensor space.

Subsequently, the resulting neural activity was split into temporal epochs. For the correct rejection trials, epochs were locked to the stimulus onset, beginning 200 ms before and extending until 800 ms after stimulus onset of the trial. Correct rejection epochs were baseline-corrected relative to the pre-stimulus-onset interval from −100 to 0 ms. For the false alarm trials, epochs were locked to the response onset, beginning 500 ms before and extending until 500 ms after response onset of the trial. False alarm epochs were baselined-corrected relative to the pre-response interval from −400 to −300 ms. Epochs with a maximum voltage greater than ±150 µV or that exceeded 5 standard deviations of the mean in terms of kurtosis and probability were excluded from further analysis. Epochs that deviated from the mean by ±50 dB in the 0-2 Hz frequency window (eye movement detection) and by +25 or −100 dB in the 20-40 Hz frequency window (muscle activity detection) were rejected as well. For the sitting condition, on average 21% of the trials were rejected based on these criteria, while for the walking condition the respective percentage was 39%. Event-related potentials (ERPs) were measured by averaging epochs for (2 motor task) x (2 cognitive task) conditions, namely four (4) experimental conditions in total. The motor task conditions were 1) sitting and 2) walking; and the cognitive task conditions were 1) correct rejections and 2) response-locked false alarms.

#### Statistical Analysis

The EEG statistical analyses were performed using the FieldTrip toolbox [86] (http://fieldtriptoolbox.org). To compare ERP waveforms between sitting and walking, paired t-tests and cluster-based permutation tests were used [87]. As stated in the Introduction, significant differences between sitting and walking were hypothesized to be found in N2 ([200, 350] ms [49, 50]) and P3 ([350, 600] ms [29, 54]) latencies and topographies during correct rejection trials, and in ERN ([−50, 100] ms [57, 58]) latencies and topographies during response-locked false alarm trials. Despite having formulated specific hypotheses about the latency and the topographies of the effects, the full set of 64 electrodes and all the epoch timepoints were included in the analyses, to explore potential effects that might have been overlooked by previous studies. By using this approach, both the hypothesis-driven and the exploratory component of this study are satisfied at once.

First, the mean walking-*minus*-sitting difference ERP waveform was obtained for each electrode and for each subject by subtracting the within-subject mean sitting ERP waveform from the corresponding mean walking ERP waveform. Next, one-sample t-tests were performed on the mean difference ERP waveforms coming from all subjects, at each electrode and timepoint. To correct for multiple comparisons, cluster-based permutation tests were performed, using the Monte Carlo method (5000 permutations, significance level of the permutation tests a = 0.05, probabilities corrected for performing two-sided tests) and the weighted cluster mass statistic [88] (cluster significance level a = 0.05, parametric cluster threshold). This procedure was performed separately for each one of the interrogated behavioral conditions of the cognitive task (correct rejection, response-locked false alarm), first for the entire cohort and, subsequently, for the IMP and nIMP subgroups. The results of the point-wise t-tests from all 64 electrodes and all timepoints were displayed as an intensity plot to efficiently summarize and facilitate the identification of the onset and general topographical distribution of walking-related changes in ERP activity. The x, y, and z axes, respectively, represent time, electrode location, and the t-statistic (indicated by a color value) at each electrode-timepoint pair.

## Results

### Group-Level Analysis

#### Cognitive Task Performance

In previous studies, correct rejection rate (CRR) and hit rate (HR) have been used to assess cognitive task performance in a Go-NoGo response inhibition task [29]. However, CRR and HR in isolation can be impacted by changes in the participant’s response criterion. The sensitivity index (d’) measures the discriminability between the Go and the NoGo conditions and is independent of the response criterion. Higher d’ scores indicate an increased ability to properly detect and respond to both Go and NoGo stimuli.

In the current cohort, d’ scores during sitting and walking were calculated and d’ differences between sitting and walking were examined using a paired t-test. Overall, d’ scores were higher during walking compared to sitting (d’sitting = 2.27 ± 1.20, d’walking = 2.50 ± 1.07; t_25_ = 2.85, p = 0.0087, Cohen’s d = 0.56) indicating better performance when participants were walking on the treadmill (Fig. 2A). This observation appears to be inconsistent with the hypothesis that there will be interference between motor and cognitive tasks (CMI) during dual-task conditions [12].

**Fig. 2.**
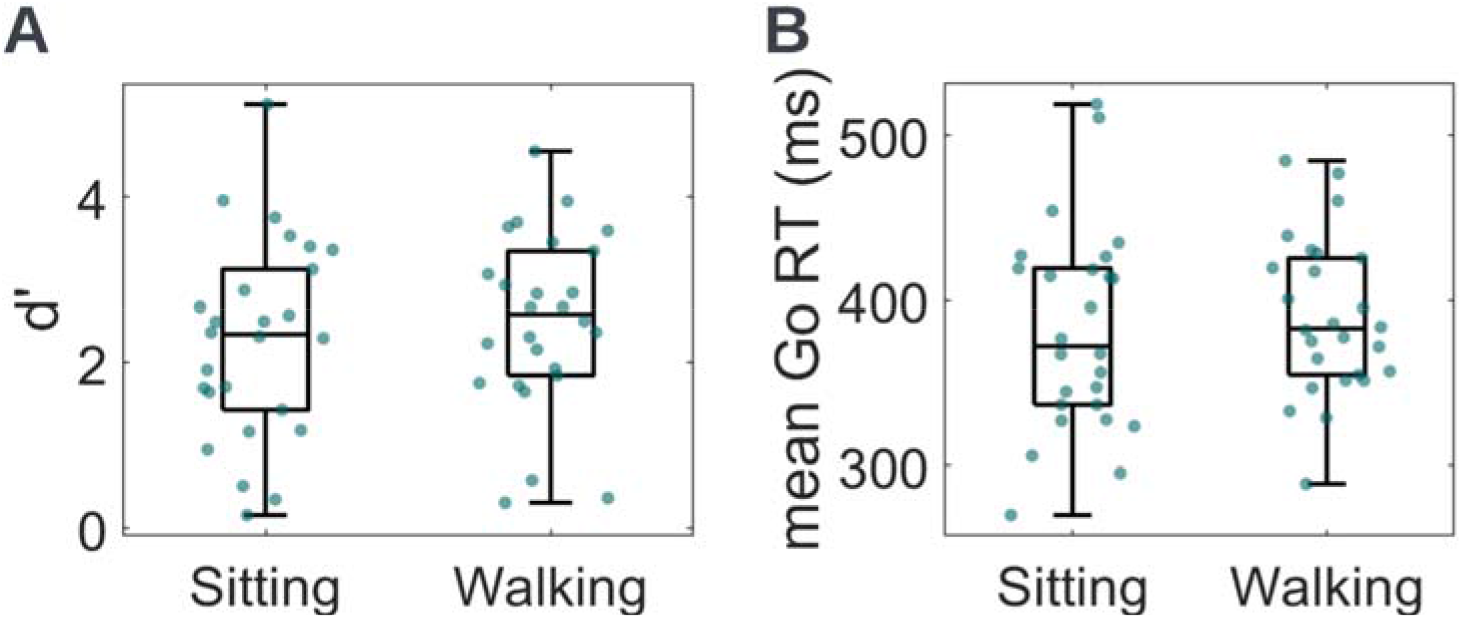
Sitting and walking **A.** d’ scores and **B.** mean Go RTs of the full young adult cohort. Dots represent individual participants. The central mark of each box indicates the median, and the bottom and top edges indicate the 25^th^ and 75^th^ percentiles, respectively. The whiskers extend to the most extreme data points not considered outliers. There were no outliers here. D’ scores during walking were higher compared to sitting, indicating better cognitive task performance during walking in young adults. No significant differences were found in mean Go RTs between sitting and walking.

Mean Go RT differences between sitting and walking were assessed using a paired t-test. No significant differences were found between sitting and walking mean Go RTs (mean RT Go sitting = 382 ± 62 ms, mean RT Go walking = 390 ± 47 ms; t_25_ = 1.36, p = 0.1862, Cohen’s d = 0.27; Fig. 2B).

Based on these data, young adults responded more accurately to task-related stimuli during walking than while sitting. This behavioral improvement was not accompanied by response speed costs (i.e., no speed-accuracy trade-off was observed).

#### Gait Kinematic Activity

Walking on a treadmill imposes a fixed walking speed, and as a result stride time variability may underestimate the impact of cognitive tasks on gait. To evaluate gait kinematics across the entire gait cycle (stance and swing phases) and compare 3D trajectories of consecutive strides, a Dynamic Time Warping (DTW) approach was used (details in Methods).

Using DTW, the variability from one stride to the next was quantified as DTW distance and, subsequently, the mean DTW distance of all stride-to-stride comparisons was extracted per participant (Fig. 3A). Mean DTW distance was calculated during both single-task (ST) and dual-task (DT) walking. One (1) participant did not have ST walking recordings and was therefore excluded from this analysis, resulting in a set of twenty-five (25) participants. Mean DTW distance differences between ST and DT walking were assessed using a Wilcoxon signed rank test. Mean DTW distances were greater during ST compared to DT walking (mean DTW distance ST = 2.44 ± 0.61 m, mean DTW distance DT = 2.21 ± 0.52 m; z = 4.02, p = 0.0001, Rosenthal’s r = 0.80). As illustrated, walking variability *decreased* when combined with the response inhibition task compared to walking in isolation (Fig. 3B).

**Fig. 3.**
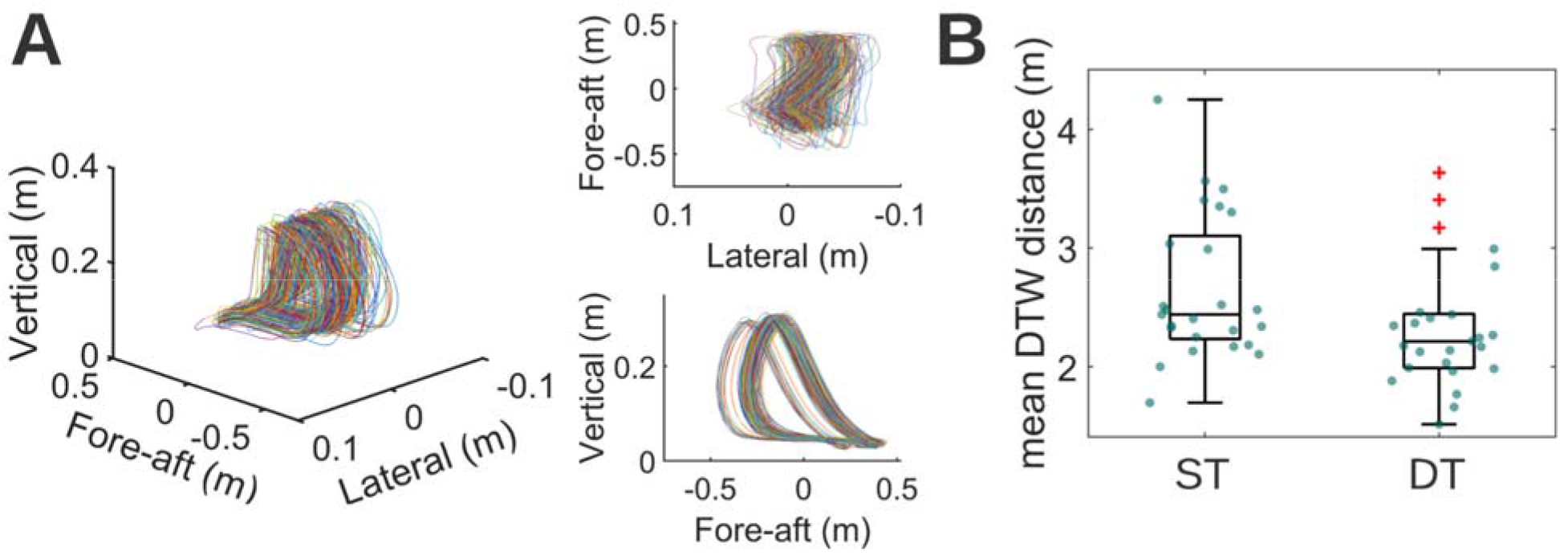
**A.** 3D representations of trajectories of a series of strides. Lateral is the dimension of movement right-and-left relative to the motion of the treadmill belt. Vertical is the dimension movement up-and-down relative to the motion of the treadmill belt. Fore-aft is the dimension of movement parallel to the motion of the treadmill belt. Perspective (left), top (upper right) and right (lower right) views are provided. Using DTW, the variability from one stride to the next was quantified as DTW distance (see Methods) and the mean DTW distance of all strideto-stride comparisons was extracted per participant. **B.** Mean DTW distance distribution during single-task walking (ST) and dual-task walking (DT); mean DTW distances of individual participants are represented as dots scattered on the boxes. Mean DTW distance was smaller during DT compared to ST walking. Red ‘+’ symbols indicate outliers.

#### EEG Activity

##### Correct Rejections

Cluster-based permutation tests were used to examine neural activity differences between sitting and walking during correct rejection trials. First, three (3) midline electrode locations – a frontocentral midline electrode (FCz), a central midline electrode (Cz) and a centroparietal midline electrode (CPz) – were plotted and inspected for differences (Fig. 4A, latency intervals of significant differences are highlighted in gray). The selection of these electrodes was based on previous studies showing that the N2 amplitude is maximal over frontocentral midline scalp and the P3 is maximal over centroparietal midline scalp [47, 89]. Reduced ERP amplitudes during walking were found at FCz and Cz during the N2 latency interval, and at the Cz and CPz during the P3 latency interval (Fig. 4A). The cluster-based permutation approach also allowed for exploring the existence of walking-related effects on ERPs in the entire electrode set and at all the epoch timepoints. The effects that this approach revealed were ERP amplitude reductions during walking 1) over frontal and frontocentral scalp (yellow in the Fig. 4B statistical clusterplot) and over parietal and occipital scalp (blue) during the N2 latency interval, 2) over left prefrontal (yellow) and over central and centroparietal scalp (blue) during the P3 latency interval and 3) over central and centroparietal scalp during latencies beyond the P3 latency interval, until the end of the epoch (blue). The topographical distribution of the average (walking-*minus*-sitting) neural activity difference during correct rejection trials is shown for selected timepoints at which this difference was found to be significant (Fig. 4C, red dots on the maps show electrodes that exhibit significant differences).

**Fig. 4.**
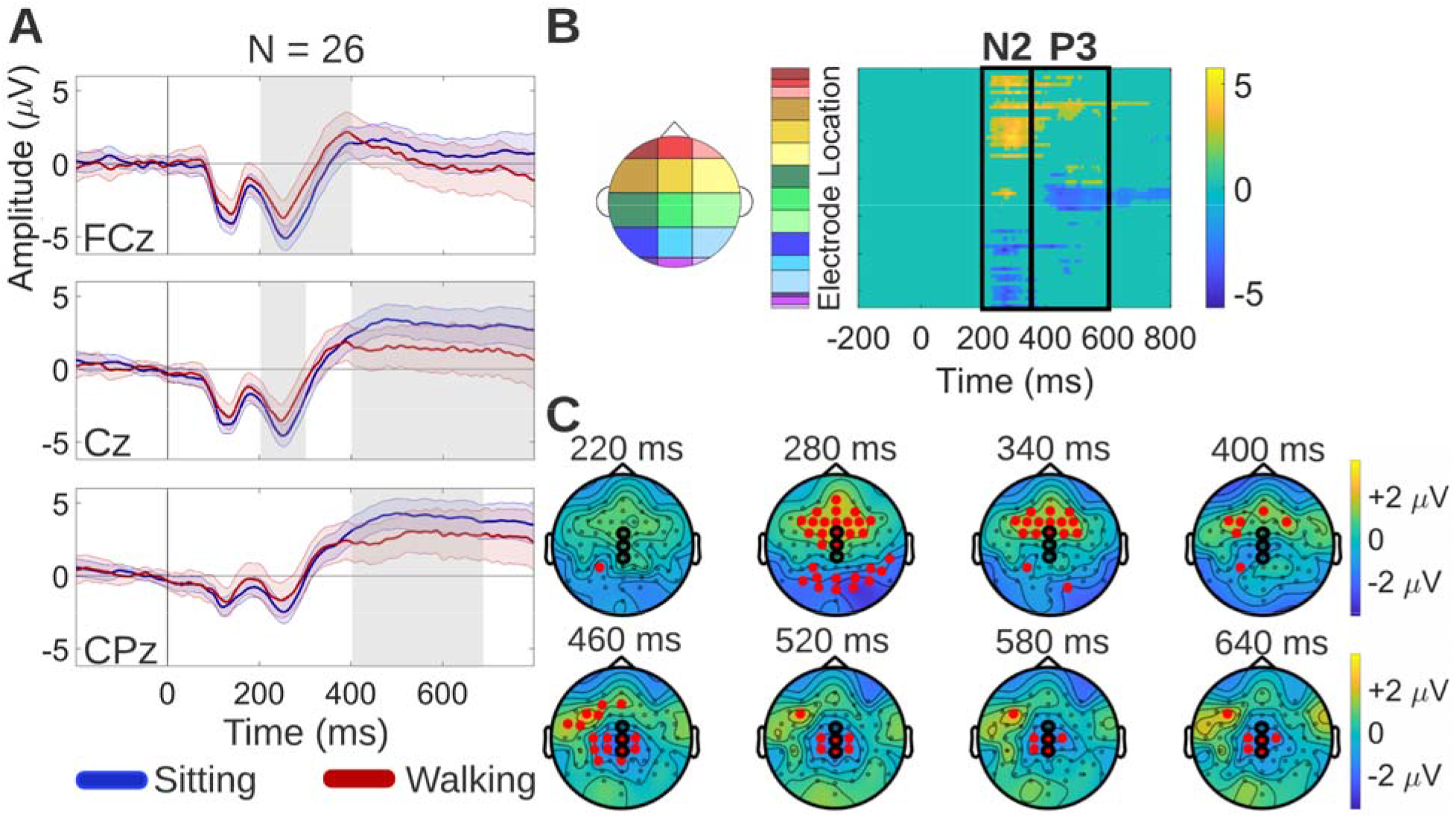
Neural activity differences between sitting and walking during correct rejection trials. **A.** Grand average sitting and walking ERP waveforms, at three midline electrode locations: frontocentral midline (FCz), central midline (Cz) and centroparietal midline (CPz) electrode. The shaded regions around the ERP waveforms indicate the Standard Error of the Mean (SEM) across participants. The latency interval of significant differences in each electrode is highlighted in gray. **B.** Spatiotemporal walking-*minus*-sitting ERP differences using cluster-based permutation tests. The statistical clusterplot shows the t-values for the electrode-timepoint pairs at which significant ERP differences between sitting and walking were found. Positive t-values (yellow) indicate that walking ERP amplitude was greater than sitting ERP amplitude. Significant differences were found 1) over frontal and frontocentral scalp (yellow) and over parietal and occipital scalp (blue) during the N2 latency interval, 2) over left prefrontal (yellow) and over central and centroparietal scalp (blue) during the P3 latency interval and 3) over central and centroparietal scalp during latencies beyond the P3 latency interval, until the end of the epoch (blue). The black rectangles indicate latencies corresponding to the N2 and P3. **C.** Topographical maps showing the average (walkingminus-sitting) neural activity difference for selected timepoints at which this difference was found to be significant. The electrodes exhibiting significant differences are depicted as red dots on the maps. The electrodes to which the ERP waveforms of panel A correspond are circled in black (vertical order matched).

##### Response-Locked False Alarms

Cluster-based permutation tests were used to examine neural activity differences between sitting and walking during response-locked false alarm trials. First, the FCz electrode was plotted and inspected for differences (Fig. 5A), since the ERN has been shown to have maximal amplitude over frontocentral midline scalp [90, 91]. Reduced ERP amplitudes during walking were found at FCz during the ERN latency interval. By exploring the entire electrode set and all the epoch timepoints, the effects that the cluster-based permutation approach revealed were frontocentral walking-related amplitude reductions during ERN latencies, as well as during latencies preceding the ERN ([−130, −50 ms] approximately, corresponding to the yellow points outside of the black rectangle in Fig. 5B statistical clusterplot). The latency and topography of these earlier, pre-ERN differences indicate reduction in pre-motor neural activity reflected by the pre-movement positivity (PMP) ERP [92, 93]. The topographical distribution of the average (walking-*minus*-sitting) neural activity difference during response-locked false alarm trials is shown for selected timepoints at which this difference was found to be significant (Fig. 5C, red dots on the maps show electrodes that exhibit significant differences).

**Fig. 5.**
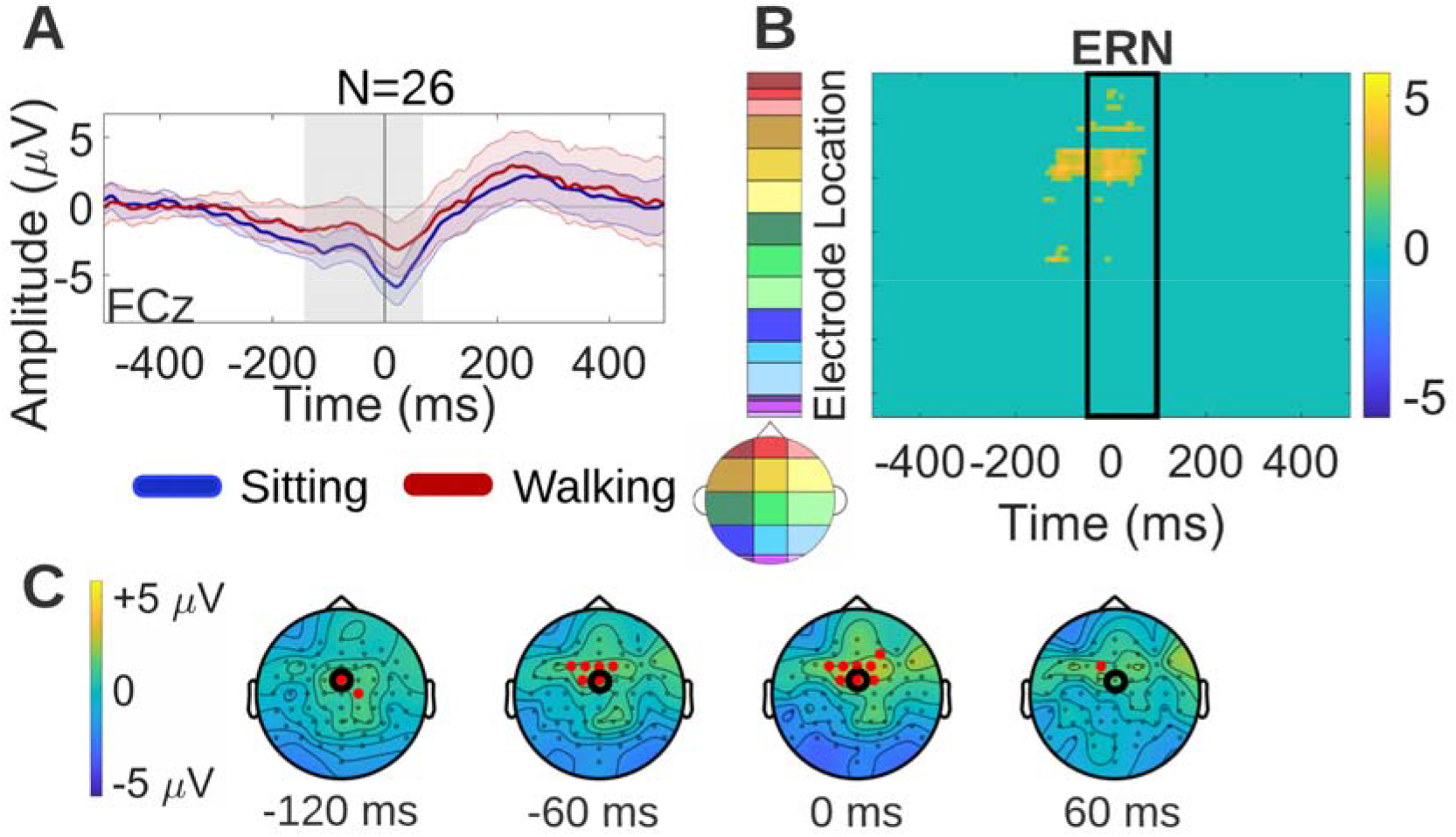
Neural activity differences between sitting and walking during response-locked false alarm trials (modeled per Fig. 4). **A.** Grand average sitting and walking ERP waveforms during response-locked false alarm trials, at a frontocentral midline electrode (FCz). **B.** Spatiotemporal walking-*minus*-sitting ERP differences using cluster-based permutation tests. Significant differences, shown in yellow in the statistical clusterplot, were found over frontocentral scalp during the ERN latency interval, and over similar frontocentral scalp during pre-ERN latencies. The latter indicated reduction in the pre-motor positivity (PMP). **C.** Topographical maps showing the average (walking-*minus*-sitting) neural activity difference for selected timepoints at which this difference was found to be significant. The electrode to which the ERP waveforms of panel A correspond is circled in black.

### Split-Group Differences Based on Cognitive Task Performance

#### Cognitive Task Performance

As shown above, mean d’ scores for the entire cohort improved when participants were walking on the treadmill. This is an indication of cognitive-motor enhancement, rather than the more typically expected cognitive-motor interference often observed in dual task paradigms. In other words, the response inhibition task appears to have gotten easier when coupled with walking in this young adult cohort. Several questions arise from this observation: Does each individual improve? Are there neural patterns that differ based on behavioral improvement versus non-improvement while walking? Are there differences in gait variability for those that improve compared to those that do not? These questions are addressed below in split-group analyses that compare those who improved their performance while walking (IMPs) and those who did not (nIMPs).

The (d’walking – d’sitting) difference was calculated for each participant and its significance was subsequently tested by determining whether it lay outside of the 95% confidence interval of the normal distribution having a mean value of zero and a standard deviation equal to that of the (d’walking – d’sitting) distribution of the entire cohort. If d’walking > d’sitting, namely the participant improved significantly during walking compared to sitting, they were classified into the IMP subgroup. If d’walking ≤ d’sitting, namely the participant did not improve significantly during walking compared to sitting, they were classified into the nIMP subgroup. In total, fourteen (14) participants were classified into the IMP subgroup and twelve (12) participants into the nIMP subgroup (8 had d’walking ≈ d’sitting, 4 had d’walking < d’sitting), as shown in Fig. 6A.

**Fig. 6.**
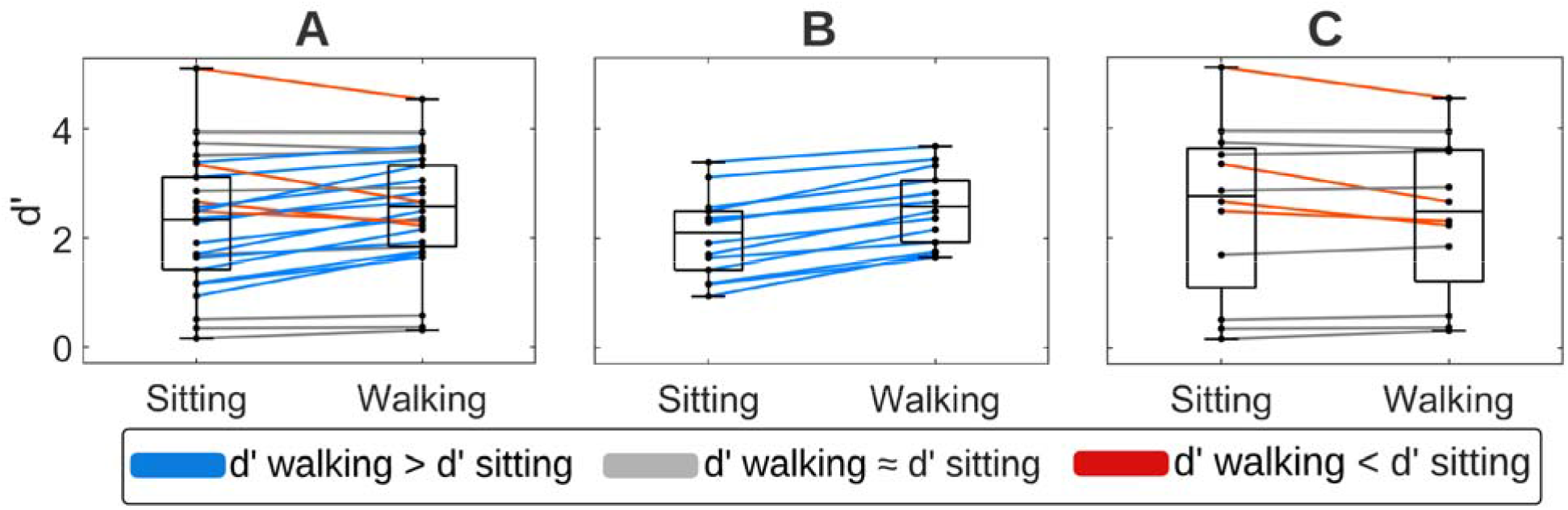
Sitting and walking d’ scores of **A.** the full young adult cohort, **B.** participants who improved during walking (IMPs), and **C.** participants who did not improve during walking (nIMPs). Each line corresponds to one participant.

In the subsequent analyses, response times, ERPs, and gait kinematic variability were contrasted between IMPs (Fig. 6B) and nIMPs (Fig. 6C).

No significant differences in average d’ scores were found between IMPs and nIMPs, as the independent samples t-test indicated (average d’ IMPs = 2.30 ± 0.70, average d’ nIMPs = 2.48 ± 1.49; t_24_ = 0.39, p = 0.7020, Cohen’s d = 0.15).

The 2×2 ANOVA assessing the effects of Group (IMPs vs nIMPs) and Motor Load (sitting vs walking) on mean RTs revealed a significant main effect of Group (F_1,24_ = 4.86, p = 0.0373, η^2^ = 0.16). This indicated that IMPs (mean RT sitting = 360 ± 54 ms, mean RT walking = 372 ± 41 ms) were overall faster than nIMPs (mean RT sitting = 407 ± 64 ms, mean RT walking = 411 ± 47 ms) to respond to image presentation. No significant effects of Motor Load (F_1,24_ = 1.65, p = 0.2108, η^2^ < 0.01) or Group/Motor Load interaction was found (F_1,24_ = 0.55, p = 0.4649, η^2^ < 0.01).

#### Gait Kinematic Activity

The effects of Group (IMPs vs nIMPs) and Cognitive Load (ST vs DT walking) on mean DTW distance were assessed by means of a 2×2 ANOVA. This ANOVA revealed a significant main effect of Group (F_1,23_ = 4.71, p = 0.0406, η^2^ = 0.14) indicating that IMPs (mean DTW distance ST = 2.46 ± 0.54 m, mean DTW distance DT = 2.08 ± 0.36 m) walked less variably than nIMPs (mean DTW distance ST = 2.82 ± 0.67 m, mean DTW distance DT = 2.59 ± 0.57 m) (Fig. 7B).

**Fig. 7.**
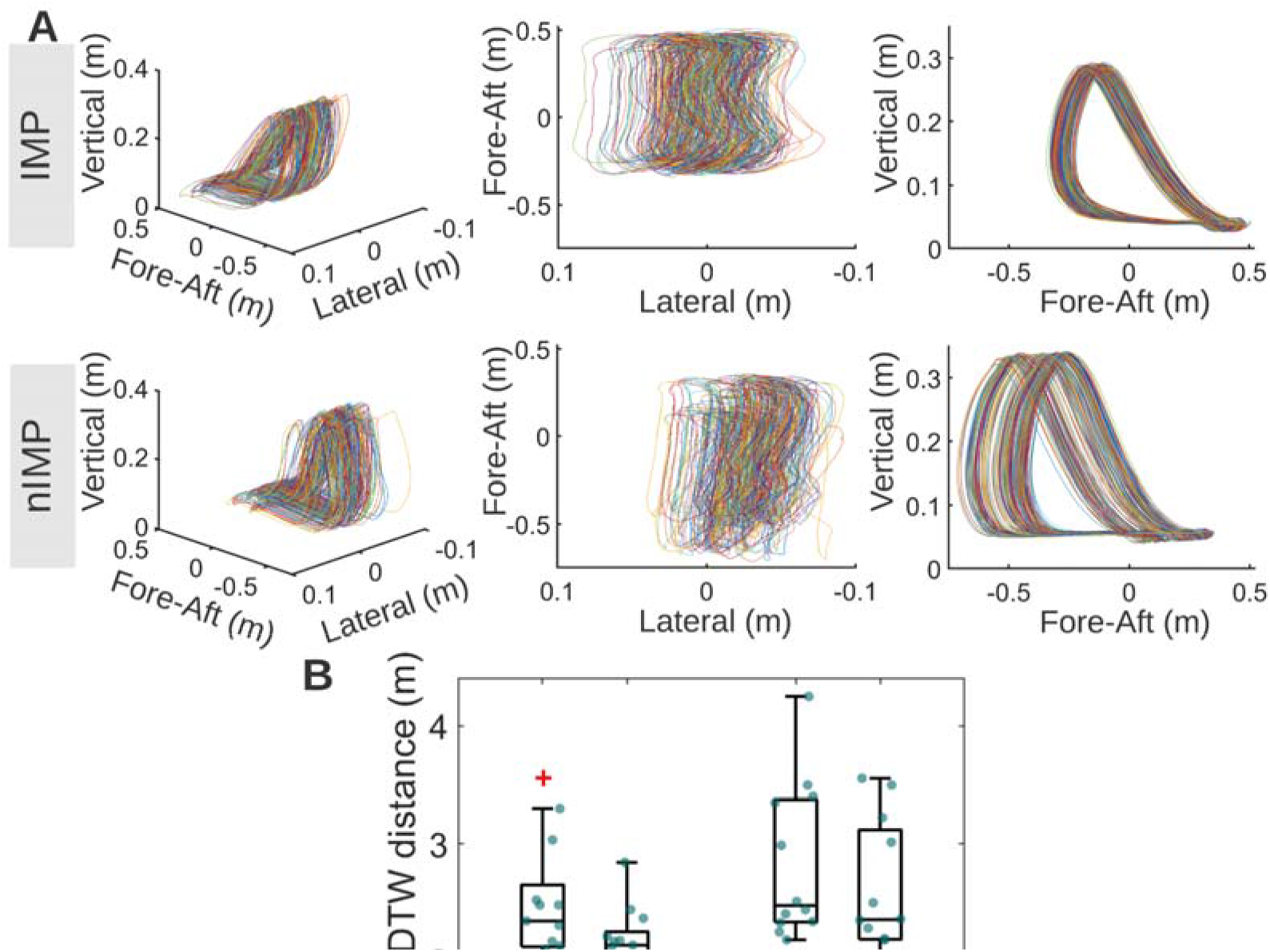
**A.** 3D representations of trajectories of a series of strides for an IMP (top) and a nIMP (bottom). For both the IMP and the nIMP, perspective (left), top (middle) and right (right) views are provided in the respective panels. **B.** Mean DTW distance distribution during single-task walking (ST) and dual-task walking (DT), for IMPs and nIMPs. Mean DTW distance was smaller in IMPs compared to nIMPs.

The significant main effect of Cognitive Load (F_1,23_ = 13.48, p = 0.0013, η^2^ = 0.07) that occurred is expected, since the same effect was tested using the Wilcoxon signed rank test as part of the group-level analysis. Within-group post-hoc t-tests showed that mean DTW distance decreased significantly during DT walking in both the IMP (t_12_ = 2.59, p = 0.0235, Cohen’s d = 0.72) and the nIMP subgroups (t_11_ = 3.35, p = 0.0064, Cohen’s d = 0.97). No significant Group/Cognitive Load interaction was found (F_1,23_ = 0.80, p = 0.3811, η^2^ < 0.01) (Fig. 7B). Of note, the one (1) participant excluded from the corresponding group-level analysis was an IMP, thus resulting in a set of thirteen (13) IMPs and twelve (12) nIMPs entered into this analysis.

Fig. 7A shows 3D representations of trajectories of a series of strides for an IMP (top) and a nIMP (bottom), to give an example of what a lower-variability series of strides (IMP) looks like compared to a higher-variability series of strides (nIMP).

#### EEG Activity

##### Correct Rejections

To compare the spatiotemporal effects of walking on neurophysiology between IMPs and nIMPs during correct rejection trials, cluster-based permutation tests were performed separately for each group to examine differences between the sitting and the walking correct rejection ERP waveform. During correct rejection trials, IMPs exhibited reduced walking ERP amplitudes over frontocentral scalp during the N2 latency interval and over left prefrontal scalp during the P3 latency interval, while nIMPs exhibited no detectable ERP differences between sitting and walking (Fig. 8B). These walking-related effects in IMPs were represented by the yellow cluster in the Fig. 8B statistical clusterplot. By comparing these walking-related effects to the corresponding group-level effects shown in the Fig. 4B statistical clusterplot, it can be observed that the yellow cluster was almost identical between IMPs and the entire cohort, indicating that IMPs maintain only the frontal/frontocentral portion of walking-related effects that were found in the overall combined cohort. Sitting and walking ERPs of IMPs and nIMPs during correct rejection trials are depicted at FCz, since this electrode belongs to the frontocentral cluster of significant effects (Fig. 8A). The topographical maps of Fig. 8C show the scalp distribution of the average (walking-*minus*-sitting) neural activity difference in IMPs and nIMPs during correct rejection trials, for selected timepoints at which this difference was found to be significant in IMPs.

**Fig. 8.**
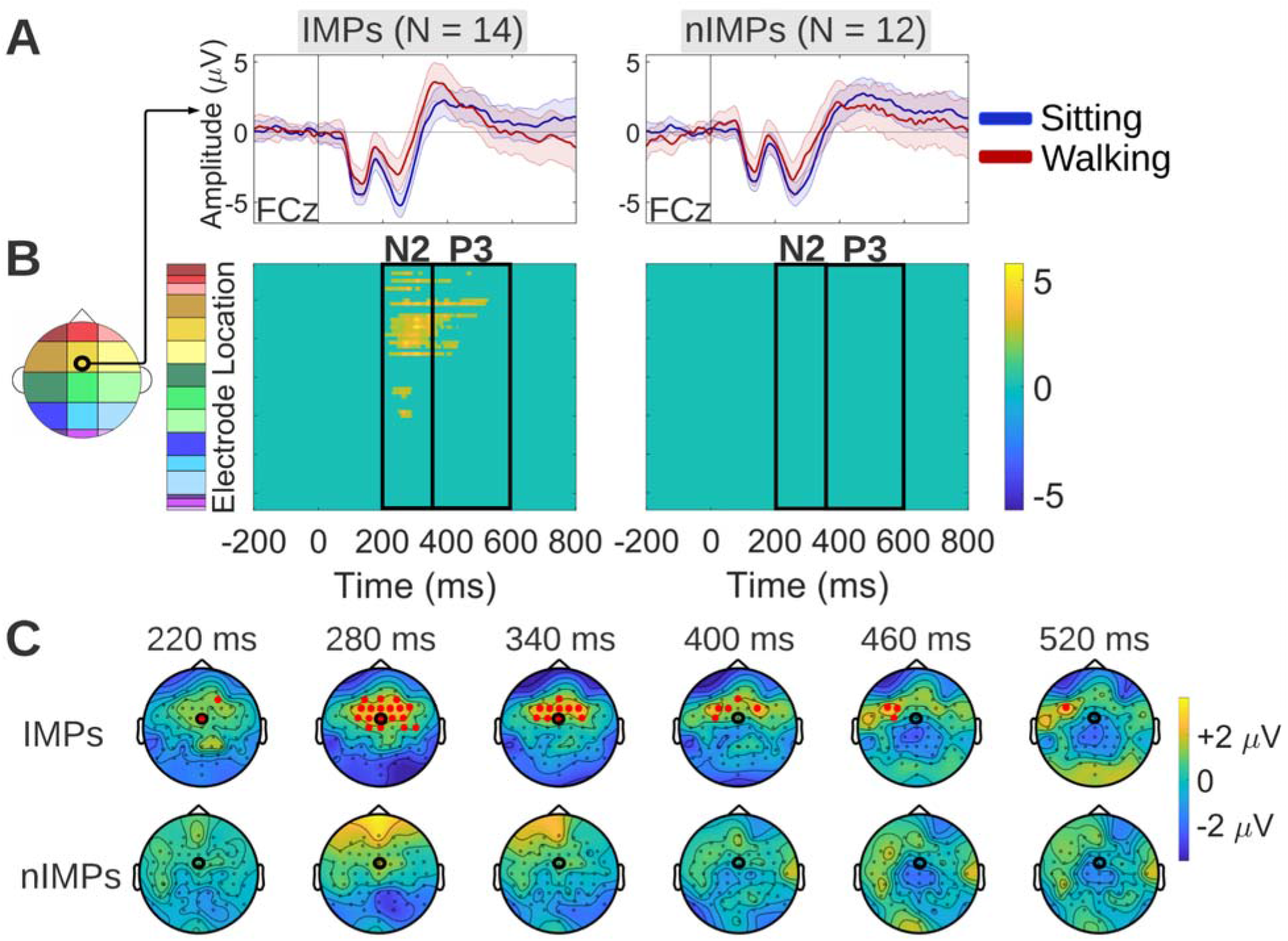
Walking-*minus*-sitting ERP differences in IMPs and nIMPs during correct rejection trials. **A.** Grand average sitting and walking ERP waveforms of IMPs (left column) and IMPs (right column), at a frontocentral midline electrode (FCz) that exhibits significant differences between the two motor load conditions for IMPs. **B.** Spatiotemporal walking-*minus*-sitting ERP differences in IMPs (left column) and IMPs (right column), using cluster-based permutation tests. Such differences were found only in IMPs, over frontocentral scalp during the N2 latency interval and left prefrontal scalp during the P3 latency interval (yellow cluster). **C.** Topographical maps showing the average (walking-*minus*-sitting) neural activity difference for selected timepoints at which this difference was found to be significant in IMPs. Maps are illustrated both for IMPs and nIMPs, for comparison purposes. The electrode to which the ERP waveforms of panel A correspond is circled in black.

##### Response-locked False Alarms

To compare the spatiotemporal effects of walking on neurophysiology between IMPs and nIMPs during response-locked false alarm trials, cluster-based permutation tests were performed separately for each group to examine differences between the sitting and walking response-locked false alarm ERP waveform. During response-locked false alarm trials, IMPs exhibited reduced walking-related ERP amplitudes over frontal/frontocentral scalp during the ERN latency interval and over central/frontocentral scalp during pre-ERN latencies (Fig. 9B). Pre-ERN amplitude reductions, which were not part of our initial hypothesis, were also encountered in the group-level analysis (Fig. 5B) where they were interpreted as reduction in the PMP. Here, even though differential effects started earlier than in the combined group (−210 vs −130 ms, approximately), they are still thought to reflect PMP amplitude reduction during walking based on their latency and topography, only more pronounced compared to the entire cohort. No detectable ERP differences between sitting and walking were found in nIMPs (Fig. 9B). The walking-related effects in IMPs are represented by the yellow cluster in the Fig. 9B statistical clusterplot. By comparing these walking-related effects to the corresponding group-level effects shown in the Fig. 5B statistical clusterplot, it can be observed that the yellow cluster manifested in IMPs was qualitatively similar to the corresponding cluster of the entire cohort, with the difference that the IMP cluster had an earlier onset (more pronounced PMP reduction during walking in IMPs) and it spread over more electrodes during the ERN latency interval (more pronounced ERN reduction during walking in IMPs). This indicates that IMPs in general maintain the walking-related effects that were found in the overall combined cohort. Sitting and walking ERPs of IMPs and nIMPs during response-locked false alarm trials are depicted at FCz, since this electrode belongs to the frontocentral cluster of significant effects (Fig. 9A). The topographical maps of Fig. 9C show the scalp distribution of the average (walking-*minus*-sitting) neural activity difference in IMPs and nIMPs during response-locked false alarm trials, for selected timepoints at which this difference was found to be significant in IMPs.

**Fig. 9.**
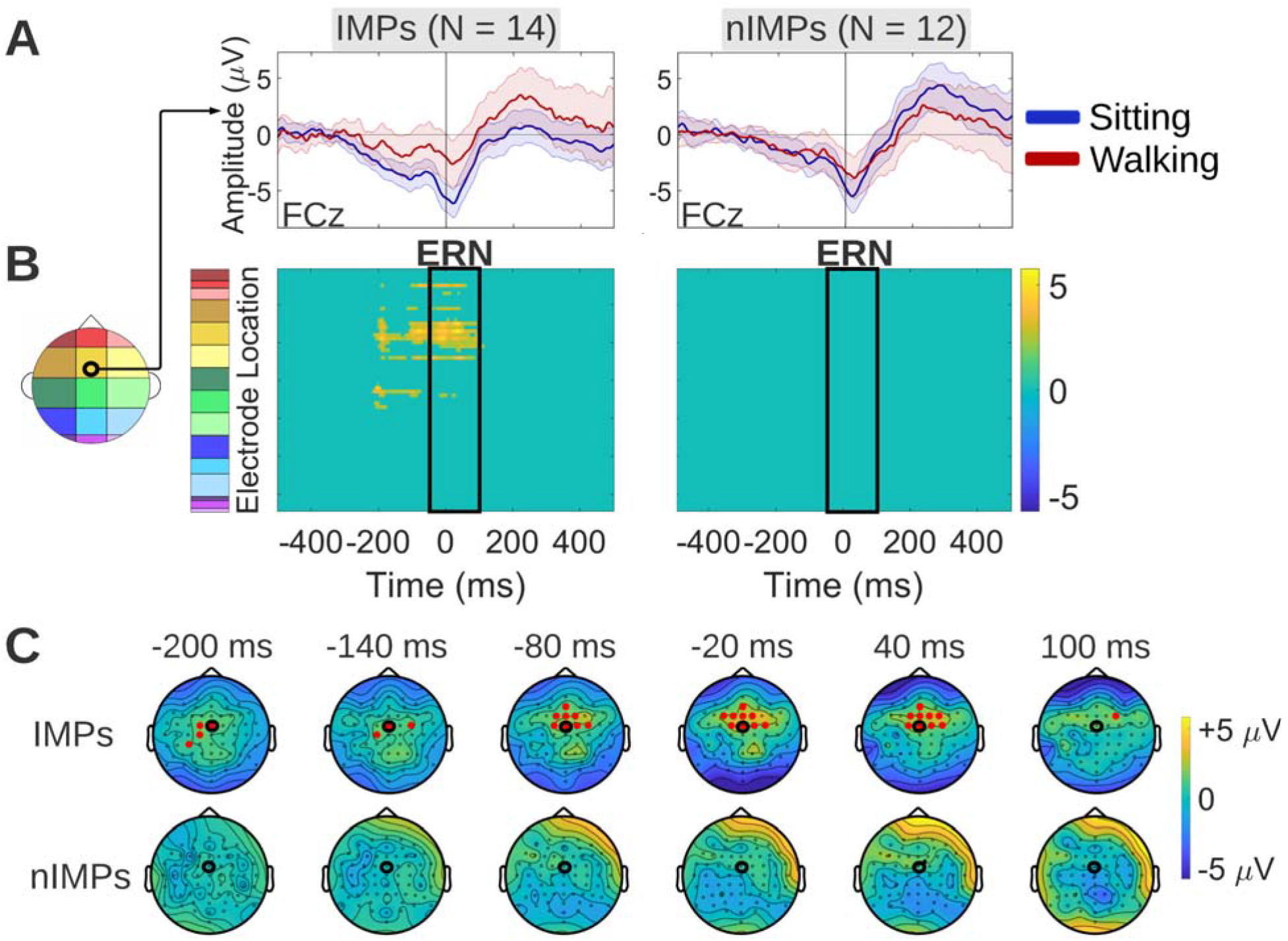
Walking-*minus*-sitting ERP differences in IMPs and nIMPs during the response-locked false alarm trials (modeled per Fig. 8). **A.** Grand average sitting and walking ERP waveforms of IMPs (left column) and IMPs (right column), at a frontocentral midline electrode (FCz) that exhibits significant differences between the two motor load conditions for IMPs. **B.** Spatiotemporal walking-*minus*-sitting ERP differences in IMPs (left column) and IMPs (right column), using cluster-based permutation tests. Such differences were found only in IMPs, over frontocentral scalp during the ERN latency interval and over central/frontocentral scalp during latencies preceding the ERN (yellow cluster). The latter indicated reduction in the PMP. **C.** Topographical maps showing the average (walking-*minus*-sitting) neural activity difference for selected timepoints at which this difference was found to be significant in IMPs. Maps are illustrated both for IMPs and nIMPs, for comparison purposes. The electrode to which the ERP waveforms of panel A correspond is circled in

## Discussion

Cognitive-Motor Interference (CMI) [14] predicts that walking will have a detrimental effect on cognitive task performance and/or on gait kinematics due to competition in neural resource allocation between the two concurrent task components. However, in the current cohort, walking actually improved performance in more than half of the participants (14 out of 26). The remaining 12 participants showed either no change in performance (n = 8) while walking, or had the expected decline (n = 4). Those who improved (IMPs) responded more quickly to Go stimuli and had significantly reduced stride-to-stride variability compared to those who did not improve (nIMPs). Behavioral improvement effects while walking were accompanied by ERP changes during both the correct rejection trials and the response-locked false alarm trials. No dual-task-related ERP differences were present in participants who did not improve performance while walking. The findings above suggest that IMPs may adjust their cognitive strategy in response to the increased task demands, and this adjustment is reflected in the ERP amplitude reductions during key processing stages of inhibitory control.

Reduction in stride-to-stride variability during dual-task walking compared to single-task walking was found both in the cohort overall and in the IMP and nIMP subgroups, suggesting that this effect may characterize young healthy adults in general, independent of cognitive task performance. A number of studies have reported findings similar to this, especially in young adults [94-99]. One possibility to account for this pattern is that shifting attention solely to motor control of walking, a largely automated motor pattern [100], increases susceptibility to endogenous or exogenous noise, which, in turn, can compromise walking performance [97, 101, 102]. In contrast, with the addition of a cognitive load, attention shifts away from walking-related motor control, enhancing automaticity and improving the consistency of the generated walking patterns [95, 98]. Indeed, a recent paper demonstrated that the addition of cognitive load, via performance of a Go-NoGo task, reduced the impact of perturbations in ongoing optic flow inputs on both gait parameters and EEG outcomes while participants were engaged in treadmill walking [103].

During correct rejection trials, ERP amplitude reductions during walking were found 1) over frontal, frontocentral, parietal and occipital scalp regions during the N2 latency interval, 2) over left prefrontal, central and centroparietal scalp regions during the P3 latency interval and 3) over central and centroparietal scalp regions during latencies beyond the P3 latency interval, until the end of the epoch (Fig. 4). From these effects, the walking-related amplitude reductions of the frontocentral N2 and the centroparietal P3 are consistent with previous literature [29]. However, the left prefrontal amplitude reductions during the P3 latency interval, along with the late central/centroparietal amplitude reductions during post-P3 latencies, have not been reported by previous studies (Fig. 4). The latter central/centroparietal effect is thought to mostly stem from differences in the EEG processing pipelines; neural activity was not highpass-filtered here thus allowing more low-frequency content to be retained. IMPs maintained the anterior (frontal/frontocentral) portion of effects found in the combined cohort, namely they exhibited ERP amplitude reductions during walking over frontal/frontocentral scalp during the N2 latency interval and in left prefrontal scalp during the P3 latency interval (Fig. 8). In contrast, nIMPs exhibited no detectable walking-related effects during correct rejection trials. (Fig. 8). Focusing on the effects found in IMPs, the N2 is thought to reflect the conflict between the two competing response tendencies (the ‘Go’ and the ‘NoGo’) and its generation has been traced to the anterior cingulate cortex (ACC), a region that plays a key role in conflict monitoring [51-53, 104, 105]. The neural activity differences between sitting and walking in IMPs during these latencies had a frontal/frontocentral topographical distribution which aligns with ACC engagement (Fig. 8). During the P3 latencies, a topographical shift of the walking-*minus*-sitting ERP differences towards left lateral prefrontal scalp regions was observed (Fig. 8). This finding too can be interpreted using the conflict monitoring hypothesis, which predicts that, once conflict between active representations is detected by the ACC, a conflict-related signal is relayed to lateral prefrontal resources, and more specifically the dorsolateral prefrontal cortex (DLPFC), where inhibitory control is implemented. In that way, the top-down attentional resources hosted by the DLPFC are activated towards implementing behavioral adjustments, to reduce conflict in ensuing trials [55]. Neuroimaging studies have shown that that increased left DLPFC activation in particular is associated with lower activation levels in conflict-related brain regions in subsequent trials of a Go-NoGo response inhibition task, thus emphasizing the central role that the left DLPFC plays in exerting top-down cognitive control [106]. Taken together, our neurophysiological findings during correct rejection trials suggest that successful inhibition in IMPs during walking is driven by modulation of the conflict monitoring (N2 stage) and the subsequent control implementation (P3 stage) neural processes compared to sitting, with such effects being absent in nIMPs.

During response-locked false alarm trials, ERP amplitude reductions during walking were found over frontocentral scalp regions during the ERN latency interval, as well as during earlier latencies preceding the ERN which were interpreted as pre-movement positivity (PMP) ERP effects (Fig. 5). IMPs exhibited a pronounced version of the walking-related ERP effects found in the combined cohort, maintaining the general topography and latency of the effects, while nIMPs exhibited no detectable effects during the response-locked false alarm trials. (Fig. 9). Focusing on the effects found in IMPs, the ERN is thought to reflect the conflict between the erroneous motor response and corrective processes, a few milliseconds following the motor response (it peaks roughly 50 ms post-response, in accord with the results shown in Fig. 9). The source of the ERN has been localized to the ACC [55, 59, 60], same as the N2, consistent with the frontocentral topographical distribution of the walking-*minus*-sitting ERP differences demonstrated in Fig. 9 during this latency interval. The main difference compared to the N2 is that here the ACC is activated and thus conflict is detected after the response instead of before [52, 55]. Therefore, the timing of the activation of the ACC, which functions as a conflict monitor, plays a critical role in determining the behavioral outcome. Regarding the earlier pre-ERN differences, the [−210, −50] ms latency interval during which they were observed has been associated with pre-motor processes reflected by the PMP [92, 93]. The PMP has been proposed to reflect the ‘go-ahead’ signal generated by the supplementary motor area (SMA) and pre-SMA to allow movement execution, consistent with the central/frontocentral topography of the walking-*minus*-sitting ERP differences detected during these latencies (Fig. 9). Summarizing the findings during response-locked false alarm trials, IMPs seem to manage an unsuccessful inhibition differently during walking compared to sitting, and this difference was pinpointed to modulation of the pre-motor neural processes preceding the erroneous motor response (PMP stage), as well as of the conflict monitoring neural processes immediately following the erroneous response (ERN stage). No such modulation was evident in nIMPs.

The CMI hypothesis proposes that cognitive task-related and gait kinematic performance decrements stem from underlying competition in the allocation of neural resources between the cognitive and the motor task. The IMP phenotype of improvement in all interrogated domains clearly contradicts the CMI hypothesis, suggesting that neural resource competition is likely absent in this subgroup, or that other factors are at play. Given that IMPs were found to alter neural activity during walking compared to sitting while nIMPs did not, these IMP-specific walking-related neural activity changes might hold the key to understanding how IMPs manage to flexibly recalibrate the underlying neural processes in order to avoid inter-task neural resource competition. It is proposed that these neural activity changes manifested by IMPs in response to motoric load increase hold promise as neural markers of cognitive flexibility.

One explanation for not observing CMI effects in IMPs is that the employed task might not have been sufficiently taxing on their cognitive resources. Walking is a relatively automatic motor task [100], so it may have sufficed to recruit certain cortical (e.g. sensorimotor) or non-cortical (e.g. subcortical, brainstem, spinal) neural networks to produce this automatic walking pattern [107-111] without interfering with the neural resources used by the response inhibition task. Also, the generally good performance of the IMP subgroup on the Go-NoGo response inhibition task suggests that this is not an especially difficult task for them. It will be interesting to observe in future work whether increasing the difficulty of the motor and/or the cognitive task will bring the cognitive resources of IMPs closer to their capacity limit, and as such, make individuals of this group manifest effects consistent with CMI. However, IMPs did not just maintain performance; they actually achieved better performance when walking. Findings such as this, although unanticipated based on CMI, have been reported and interpreted before. There is evidence that moderate exercise like walking can enhance sustained attention and facilitate cognitive task performance [112-114]. This facilitatory effect has been explained using neurotransmitter models, proposing that moderate exercise induces an increase in catecholamine levels, which, in turn, boosts the signal-to-noise ratio during processing by prefrontal attentional systems [115, 116]. This hypothesis aligns with the present findings, since the observed modulation in lateral prefrontal neural activity in IMPs coexisted with reduced conflict-related ERP amplitudes (N2 in correct rejections and ERN in response-locked false alarms) during walking, as shown in Figs. 8 and 9. Therefore, this reduction in inhibitory conflict that IMPs manifest during walking might stem from a more active and/or efficient engagement of top-down attentional resources, which presumably induces a shift to a more proactive cognitive strategy and hence promotes better anticipation of the rare ‘NoGo’ events.

A limitation of this study is that nIMPs were more variable in terms of d’ scores than IMPs, both for sitting and walking. Based on Fig. 6, while IMPs all lie within a range of medium d’ performance, nIMPs were scattered over the full d’ range, encompassing low, medium and high performers. As part of future research, we aim to collect more nIMP datasets in order to compare IMPs and nIMPs having d’ scores within similar ranges. Despite this limitation, the neural signatures of improvement that this study yielded hold significant potential as cognitive flexibility markers which could potentially be translated to older neurotypical or patient populations to assess and quantify age-related or neurodegeneration-related cognitive decline, respectively. As a first step in this direction, we aim to expand the methodology employed here to older neurotypical adults to test its efficacy in distinguishing ‘super-agers’ from older adults that exhibit normal or aggravated age-related cognitive decline [117].

## Conclusions

This study examined differences in neural activity, stride-to-stride variability and response speed between 1) young adults who improved in terms of response accuracy during walking compared to sitting (IMPs) and 2) young adults who did not improve (nIMPs) under the same conditions. This split of the young adult cohort was motivated by findings at the piloting stage of the study, where 3 out of 5 young adults performed better during walking compared to sitting, conflicting with the CMI hypothesis. During correct rejection trials, ERP amplitude reductions were found during walking in IMPs, specifically over frontocentral scalp regions during N2 latencies and over left prefrontal scalp regions during P3 latencies. Also, during response-locked false alarm trials, IMPs exhibited reduced ERP amplitudes while walking over frontocentral scalp regions, during Event-Related Negativity (ERN) and pre-ERN latencies. No detectable differences were found in the neural activity of nIMPs between sitting and walking, neither during correct rejection trials nor during response-locked false alarm trials. The present findings indicate that IMPs can flexibly modulate frontal brain activity during walking during key stages of inhibitory control (conflict monitoring for N2 and ERN, control implementation for P3, pre-motor for pre-ERN), something that nIMPs do not seem to do. Combining these neurophysiological findings with findings of faster responses and less stride-to-stride variability in IMPs, these neural activity differences were interpreted as neural signatures of behavioral improvement during dual-task walking. Future research can test the potential of these neural signatures as markers for assessing cognitive flexibility in populations where it tends to get compromised, for example older adults who either age normally or have been diagnosed with neurodegenerative diseases, such as Parkinson’s or Alzheimer’s disease.

## Abbreviations List

CMI: Cognitive Motor Interference
DT: Dual Tasking
ST: Single Tasking
RT: Response Time
EEG: Electroencephalography
ERP: Event-Related Potential
ERN: Event-Related Negativity
PMP: Pre-Motor Positivity
MoBI: Mobile Brain-Body Imaging
3D: Three-Dimensional
DTW: Dynamic Time Warping
ACC: Anterior Cingulate Cortex
DLPFC: Dorsolateral Prefrontal Cortex
IMPs: participants who improved in terms of cognitive task performance
nIMPs: participants who did not improve in terms of cognitive task performance

## Acknowledgements

We would like to thank each of the participants that enrolled in the study. Special thanks to George Kassis, Emma Mantel, Nicholas Abraham, Soma Mizobuchi and Suzan Hoffman for their help with this study.

## Data Sharing Statement

Data from this study will be made available through a public repository (e.g. Figshare) upon publication of this paper, and the authors will work with the editorial office during production to incorporate appropriate links. Custom code from this study will be made available on GitHub (https://github.com/CNL-R) upon publication of this paper.

