## Supplementary Materials for "Young adults who improve performance during dual-task walking show more flexible reallocation of cognitive resources: A Mobile Brain-Body Imaging (MoBI) study"

**Supplementary Material**

### **Pilot Data**

D’ scores

Five (5) participants ages 18-25 (1 female, 4 male) participated in the piloting of the study. All participants reported no diagnosed neurological conditions, no recent head injuries, and normal or corrected-to-normal vision. The d’ scores of the 5 pilot participants on the employed Go-NoGo response inhibition task, during sitting and walking, are listed in Supplementary Table 1. Three (3) out of 5 pilot participants exhibited greater d’ scores, therefore improved response accuracy, during walking compared to sitting. The d’ scores of these 3 participants are highlighted in bold in the table.

Supplementary Table 1. Sitting and walking d’ scores of the five (5) pilot participants. The scores of those who exhibited improved d’ performance during walking compared to sitting are highlighted in bold.

|  | Sitting | Walking |
| --- | --- | --- |
| Participant 1 (P1) | **0.58** | **0.93** |
| Participant 2 (P2) | **3.34** | **3.89** |
| Participant 3 (P3) | 3.84 | 3.30 |
| Participant 4 (P4) | **3.43** | **4.35** |
| Participant 5 (P5) | 4.04 | 3.31 |

Number of trials per condition

Supplementary Table 2 demonstrates the number of trials per behavioral condition of the Go-NoGo task, both for the walking and for the sitting condition, for each one of the 5 pilot participants. Of note, at the piloting stage, an experimental session consisted of 11 blocks: 1 training block at the beginning, 5 sitting blocks, 5 walking blocks. The only exception was participant 3 (P3) for whom only two (2) sitting and four (4) walking blocks were recorded, plus training. For the actual cohort, the number of experimental blocks was increased by adding 2 walking blocks (7 in total), 2 sitting blocks (7 in total) and 1 single-task walking block. The block order was pseudorandomized both at the piloting and at the actual cohort stage.

Supplementary Table 2. Number (#) of correct rejection trials (CR), false alarm trials (FA), hit trials (H) and miss trials (M), during sitting and walking, for each one of the 5 pilot participants.

|  | Sitting | | | | Walking | | | |
| --- | --- | --- | --- | --- | --- | --- | --- | --- |
|  | # CR | # FA | # H | # M | # CR | # FA | # H | # M |
| P1 | 29 | 112 | 961 | 84 | 45 | 98 | 963 | 82 |
| P2 | 143 | 35 | 1223 | 8 | 135 | 20 | 1042 | 3 |
| P3 | 49 | 13 | 418 | 0 | 75 | 49 | 835 | 1 |
| P4 | 109 | 46 | 1043 | 2 | 132 | 23 | 1045 | 0 |
| P5 | 139 | 13 | 1041 | 4 | 132 | 20 | 1030 | 15 |
